# Evidence for direct control of neurovascular function by circulating platelets in healthy older adults

**DOI:** 10.1101/2024.05.31.596788

**Authors:** Gabriella M.K. Rossetti, Joanne L. Dunster, Aamir Sohail, Brendan Williams, Kiera M. Cox, Suzannah Rawlings, Elysia Jewett, Eleanor Benford, Julie A. Lovegrove, Jonathan M. Gibbins, Anastasia Christakou

**Author notes:** **Footnotes** This manuscript was first published as a preprint: Rossetti GMK, Dunster JL. Sohail A, Williams B, Cox KM, Jewett E, Benford E, Lovegrove JA, Gibbins JM, Christakou A. (2024). Evidence for direct control of neurovascular function by circulating platelets in healthy older adults. bioRxiv. https://www.biorxiv.org/content/10.1101/2024.05.31.596788v1.

## Abstract

Platelets play a vital role in preventing haemorrhage through haemostasis, but complications arise when platelets become overly reactive, leading to pathophysiology such as athero-thrombosis. Elevated haemostatic markers are linked to dementia and predict its onset in long-term studies. Despite epidemiological evidence, the mechanism linking haemostasis with early brain pathophysiology remains unclear. Here, we aimed to determine whether a mechanistic association exists between platelet function and neurovascular function in 52 healthy mid- to older-age adults. To do this we combined, for the first time, magnetic resonance imaging (MRI) of neurovascular function, peripheral vascular physiology, and in vitro platelet assaying. We show a direct association between platelet reactivity and neurovascular function that is both independent of vascular reactivity and mechanistically specific: Distinct platelet signalling mechanisms (Adenosine 5’-diphosphate, Collagen-Related Peptide, Thrombin Receptor Activator Peptide 6) were directly associated with different physiological components of the haemodynamic response to neural (visual) stimulation (full-width half-maximum, time to peak, area under the curve), an association that was not mediated by peripheral vascular effects. This finding challenges the previous belief that systemic vascular health determines the vascular component of neurovascular function, highlighting a specific link between circulating platelets and the neurovascular unit. Since altered neurovascular function marks the initial stages of neurodegenerative pathophysiology, understanding this novel association becomes now imperative, with the potential to lead to a significant advancement in our comprehension of early dementia pathophysiology.

**Key points summary:**

- Haemostasis (platelet function) has been linked to the early stages of dementia, but the precise mechanisms are not well understood.
- This study asks whether a causal mechanism exists through athero-thrombotic effects on the vasculature which can in turn affect brain health, or through direct platelet effects on brain physiology.
- Here we show that elevated platelet reactivity is associated with blunted (delayed, shorter, and smaller) blood flow responses to neural activation in healthy middle-aged and older adults.
- However, the association between platelet reactivity and neurovascular function was not mediated by systemic vascular reactivity.
- This finding challenges the previous belief that systemic vascular health determines the vascular component of neurovascular function, highlighting a specific link between circulating platelets and the neurovascular unit in early dementia pathophysiology.

## Introduction

Dementia is a major cause of disability and dependency among the elderly, affecting over 55 million people worldwide and with an annual economic cost of 1.3 trillion US dollars(*1*). Clinical evidence suggests a link between dementia and cardiovascular health, including the regulation of platelets. While platelets play a vital role in our physiological well-being by preventing hemorrhage through hemostasis (blood clotting), pathological conditions arise when platelets become overly reactive, leading to complications such as athero-thrombosis(*2*). Understanding the potential implications of platelet function for dementia may elucidate preventive measures and therapeutic interventions aimed at addressing this pressing public health concern.

Hemostatic markers including platelet reactivity and pro-clotting factor concentrations (e.g. Von Willebrand factor), are elevated in individuals with vascular dementia(*3*). This association between platelet function and dementia extends beyond vascular dementia, as demonstrated in the Framingham Heart Study, where platelet aggregation response to adenosine 5’-diphosphate (ADP) at middle-age independently predicted both all-cause dementia at 20-year follow-up(*4*). In fact, platelet reactivity outweighed other commonly acknowledged risk factors, including education, LDL cholesterol, and BMI. On the basis of such associations, the use of anti-thrombotic medications has been proposed as a potential approach to slow the progression of the disease in individuals with dementia(*5*, *6*). However, the underlying physiological mechanisms require further elucidation before appropriate interventions can be effectively implemented(*7*). Taken together, this suggests that targeted prophylactic intervention on platelet function holds promise for prolonging neurocognitive health in ageing. However, evidence is limited to clinical dementia states where it is impossible to establish mechanistic links or differentiate cause and consequence.

A possible explanation of the observed relationship between platelet reactivity and dementia may be through systemic vascular effects. Indeed, dysregulation of the hemostatic system, leading to unwarranted platelet activation and subsequent thrombosis, forms an early and central aspect of cardiovascular disease pathophysiology(*8*). At the same time, subclinical vascular dysregulation emerges as an early pathological event in the development of dementia, manifesting years before detectable amyloid beta or tau abnormalities(*9*). Further, cardiovascular and metabolic health conditions, including hypertension(*4*), heart disease(*10*), and diabetes(*11*), are risk factors for mild cognitive impairment (MCI), vascular dementia, and Alzheimer’s disease(*12*).

Alternatively, platelets may directly interact with the ‘neurovascular unit,’ a complex network of neurons, glial cells, and blood vessels that regulate cerebral blood flow in response to neural activity. Supporting this, high plasma fibrinogen levels (a marker of hemostasis) have been associated with a progressive decline in cognitive abilities among the elderly, even after accounting for cardiovascular morbidity and associated risk factors(*13*), suggesting a potential direct link between platelet activity and brain function.

Notably, the signaling mechanisms involved in the regulation of platelets share similarities with those governing vascular tone. For instance, nitric oxide (NO) and prostanoids are vasodilators that are essential for synaptically-driven initiation of the hemodynamic response(*14*), and also act as potent inhibitors of platelet aggregation(*15*, *16*). Considering these interconnected factors, platelets may provide a valuable blood biomarker of cardiovascular-associated risk for dementia, facilitating the development of targeted and effective prevention or treatment strategies.

Despite growing evidence for associations between platelet function and dementia, the current evidence is dependent on disease state comparisons (i.e. dementia versus healthy), behavioral cognitive effects (e.g. nonverbal reasoning in Rafnsson(*13*)), and beta-amyloid deposition. These disease processes occur later than the dysregulation of blood vessels(*9*, *17*), and potentially much later than neurovascular dysfunction, which is proposed to be the first point of failure in the pathophysiological cascade(*14*, *18*). Despite its importance, assessment of neurovascular function in humans is complex and thus rarely performed. However, a growing number of studies have demonstrated consistent effects by estimating the model parameters of the Hemodynamic Response Function (HRF) using Blood Oxygen Level Dependent (BOLD) functional Magnetic Resonance Imaging (fMRI). Specifically, older age(*19*), diabetes(*20*, *21*), and subjective cognitive decline(*22*) are associated with altered hemodynamic responses to neural activity that are smaller in magnitude (peak height), delayed (longer time to peak; TTP), and of shorter duration (smaller full width half maximum; FWHM) compared to controls. These reduced hemodynamic responses likely reflect problems matching the vascular response to the energy demand in the active brain region. The unexplored interactions between platelets and neurovascular function could be crucial for developing targeted preventive strategies against dementia, potentially transforming global health outcomes.

We therefore sought to investigate whether a relationship exists between platelet reactivity and neurovascular function in the general population of middle- to older-aged healthy adults. Platelet reactivity was assessed using Platelet Reactivity Multiparameter Phenotyping analysis and neurovascular function was assessed by estimating hemodynamic responses (HRFs) to a fixed visual stimulus. We incorporated experimental manipulations that targeted peripheral vascular reactivity and cerebrovascular reactivity to determine whether any association could be attributed to systemic vascular effects, or direct interactions between platelets and the neurovascular unit.

## Results

### Platelet reactivity was associated with specific components of neurovascular function

#### Platelet reactivity multiparameter phenotyping revealed associations between platelet reactivity and neurovascular function

Platelet reactivity phenotyping facilitates the investigation of the underlying causes and consequences of variations in platelet function, thereby advancing progress towards precision medicine(*23*). In this study we used multiparameter phenotyping to estimate latent platelet reactivity components for each participant in order to stratify the sample (Fig. 1) and investigate association with aspects of neurovascular function (Fig. 3).

**Fig. 1.**
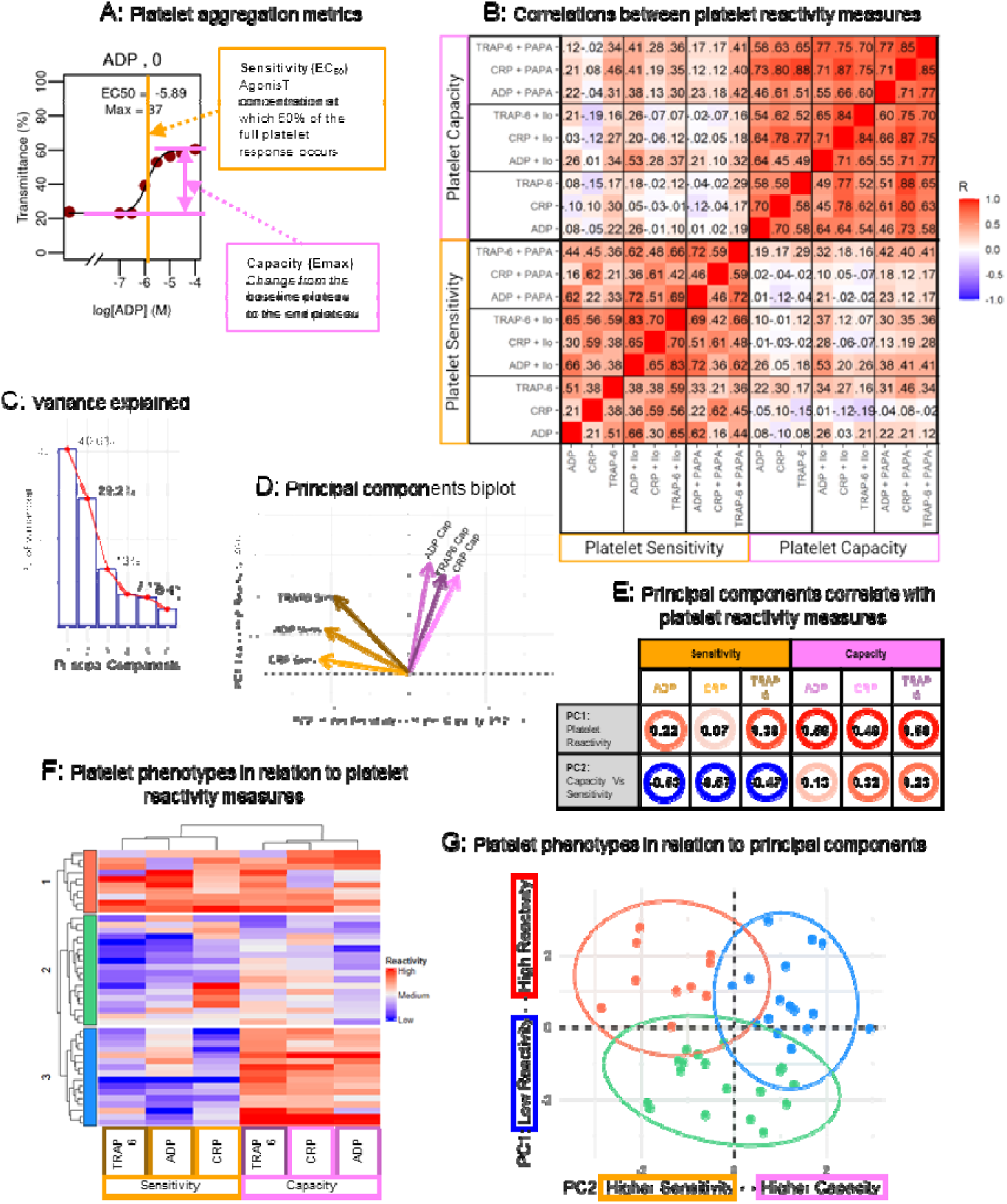
Platelet reactivity methods: Multiparameter phenotyping. Platelet reactivity was assessed by plate-based aggregometry (PBA) assay. (A) Concentration dose-response curves were analyzed to extract the metrics Sensitivity (EC50) and Capacity (Emax). For illustrative purposes, Sensitivity variables are highlighted in orange, and Capacity variables are highlighted in pink throughout the panels. throughout Platelet reactivity (Sensitivity and Capacity) was tested in response to agonists Adenosine 5’-diphosphate (ADP), Collagen-Related Peptide (CRP) and Thrombin Receptor Activator Peptide 6 (TRAP-6). Reactivity was measured to each agonist alone, and in the combined presence of each agonist with the following inhibitors; PAPA-NONOate (PAPA; a Nitric Oxide donor) and Iloprost (Ilo; synthetic prostacyclin). (B) Correlations between platelet reactivity measures confirmed Sensitivity and Capacity are distinct concepts. Values displayed are Spearman’s rho, with color reflecting the strength of the relationship. Latent platelet reactivity characteristics were identified using principal component analysis (PCA) on the agonist-only conditions (without the presence of PAPA or Ilo inhibition). (C) The first two principal components accounted for most of the variance (cumulative variance = 69.8%) and were interpreted using (D) PCA biplot and (E) patterns of correlations with the original platelet reactivity measures. Agglomerative hierarchical clustering identified three distinct platelet phenotype groups (G1, N = 10; G2, N = 18; G3, N = 16) which were reviewed in respect to (F) the original platelet reactivity measures, and (G) the principal components from the PCA.

The latent platelet reactivity components and phenotype groups were then used to investigate the effect of platelet reactivity on neurovascular function assessed by estimating local changes in blood flow in response to neural activity in the primary visual cortex (V1). The model parameters were then extracted from individual participant Hemodynamic Response Functions (HRFs; Fig. 2). Specifically, we calculated the full-width half-maximum (FWHM) which reflects the duration of the neurovascular response, time to peak (TTP) which reflects the speed of the initial neurovascular response, area under the curve (AUC) which reflects the total size of the HRF, and peak height (peak) which reflects the maximum magnitude of the neurovascular response.

**Fig. 2.**
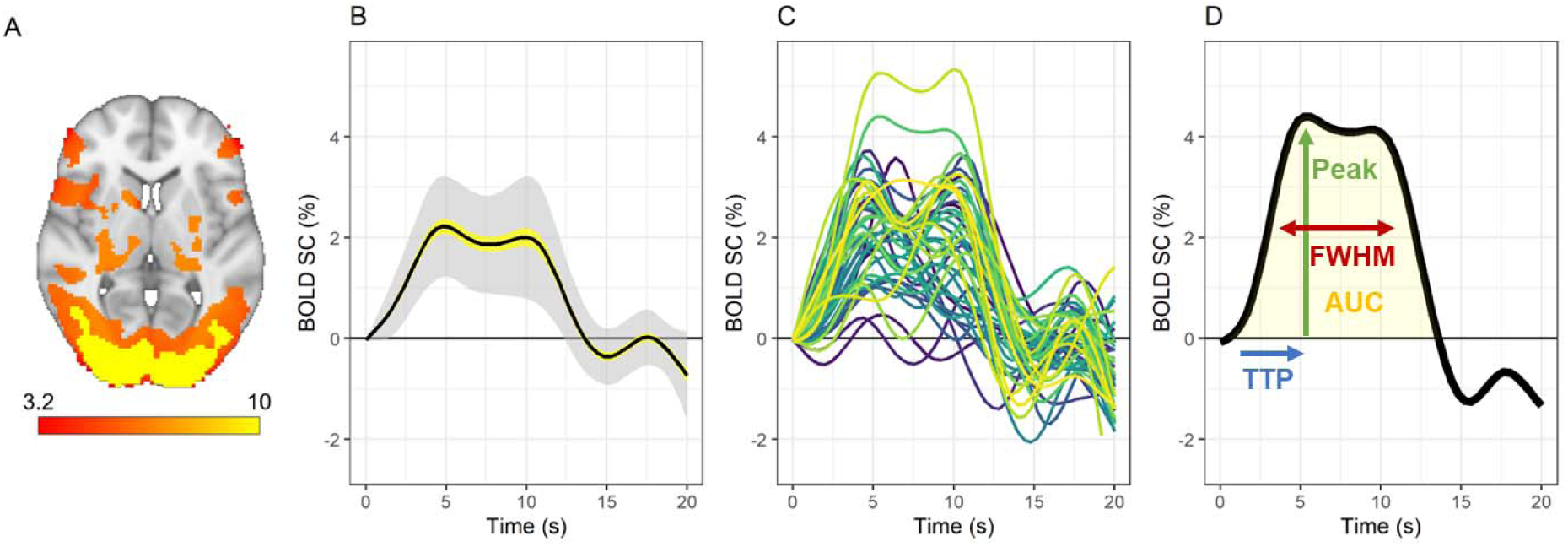
Visual stimulation evoked individualized hemodynamic response functions (HRFs). Visual stimulation using flashing checkerboard was confirmed to activate (trigger a blood flow response) in the primary visual cortex (V1). This is demonstrated by (A) Group-level activation map (color = z score); (B) Group-level average (± SE in yellow shading, ± SD in light grey shading) hemodynamic response function (HRF), and (C) Subject-specific individual HRFs (colors differentiate participants). (D) Subject-specific individual HRFs were then characterized by the parameters; full-width half-maximum (FWHM), area under the curve (AUC), time to peak (TTP), and peak height (peak).

Platelet reactivity was associated with the duration (HRF FWHM) and size (HRF AUC) of the blood flow response in the V1. Specifically, higher platelet reactivity (low -> high PC1, 40.6% variance in platelet responses, Fig. 1) was associated with shorter FWHM (R = -0.45, p = 0.006, Fig. 3A), and smaller AUC (R = -0.41, p = 0.015, Fig. 3B).

**Fig. 3.**
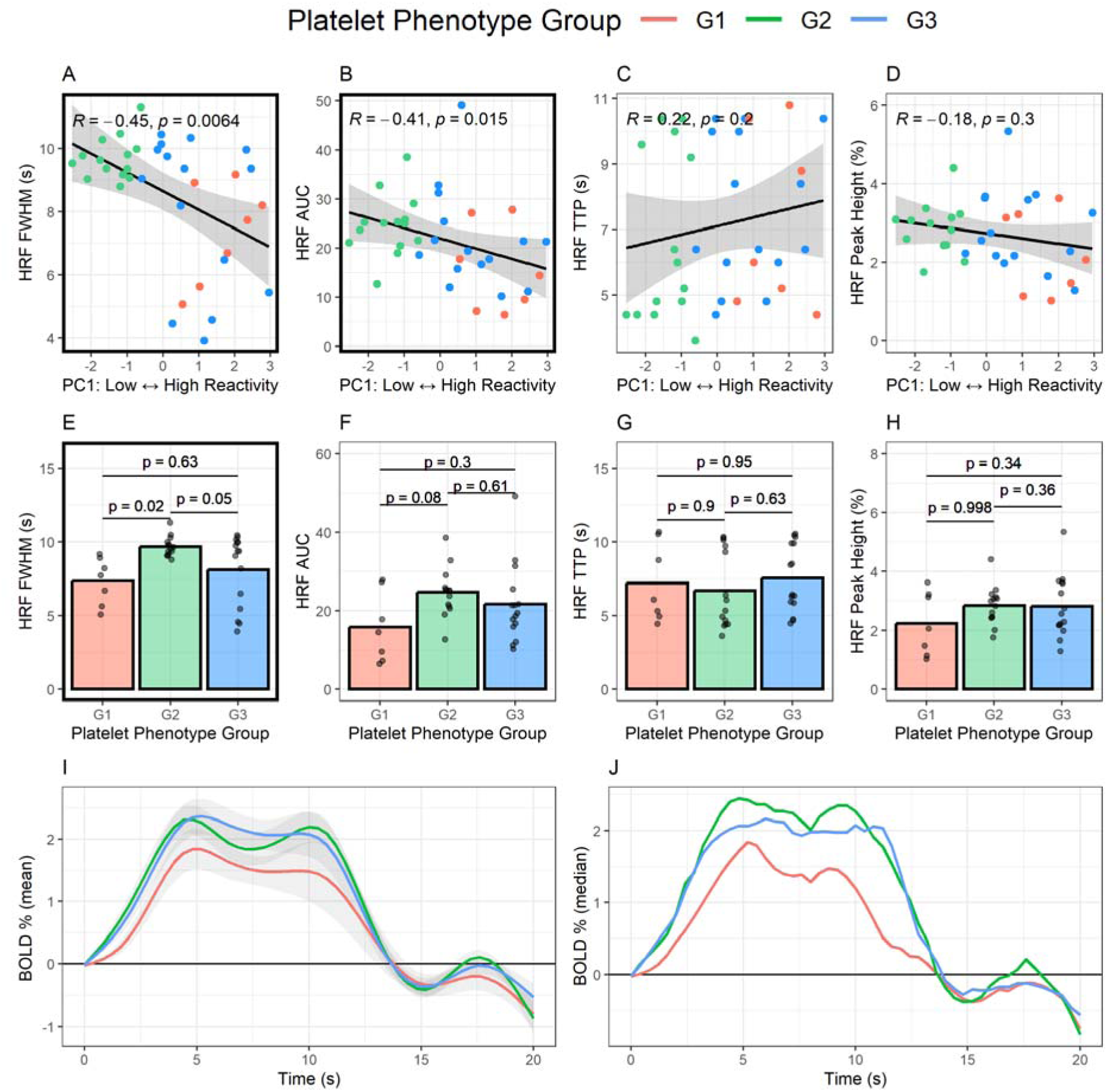
Platelet reactivity multiparameter phenotyping revealed associations between platelet reactivity and neurovascular function. For all panels colour indicates platelet phenotype group (red [G1], N = 10; green [G2], N = 18; blue [G3], N = 16). Higher platelet reactivity was associated with (A) shorter neurovascular response in the primary visual cortex (V1) measured by hemodynamic response function (HRF) full-width half maximum (FWHM) and (B) smaller neurovascular response measured by HRF area under the curve (AUC). Platelet reactivity was not associated with (C) the speed of the initial neurovascular response (time to peak; TTP) or (D) the maximum magnitude of the neurovascular response (peak height). (E) One of the high platelet reactivity groups (Group 1, see Fig 1D and Fig 1E) had a shorter FWHM compared to the low reactivity group (Group 2). There were no differences between platelet phenotype groups in (F) AUC), (G) TTP or (H) peak height. The (I) Mean (± standard deviation) and (J) Median hemodynamic response function (HRF) values for each platelet phenotype.

Participants were then grouped into subsets (phenotypes) characterized by similar patterns in platelet responses. Clustering the data revealed distinct and robust platelet phenotype groups (Fig. 1F and 1G, see Methods: Assessment of platelet reactivity and multicomponent phenotyping). These platelet phenotype groups were able to differentiate neurovascular function even in this healthy population. Specifically, the duration of the neurovascular response in the V1 (HRF FWHM) was significantly different between the platelet phenotype groups (main effect F = 5.032, p = 0.01, Fig. 3E), with pairwise group differences revealing a shorter FWHM in one of the high platelet reactivity groups (Group 1) compared to the low reactivity group (Group 2; Δ-2.34, [-0.34 – -4.34], p = 0.02, Fig. 3D). There were no significant differences between any other platelet phenotype groups (Group 1 – Group 3, Δ-0.75, [-2.73 – 1.23], p = 0.63; Group 2 – Group 3; Δ1.59, [-0.02 – 3.19], p = 0.05).

Multiparameter phenotyping did not reveal any associations between platelet reactivity and the speed of the initial neurovascular response in the V1 (HRF TTP) or the maximum magnitude of the neurovascular response (peak height). Specifically, platelet reactivity (low -> high, PC1, Fig. 1C – 1E) was not associated with TTP (R = 0.21, p = 0.20, Fig. 3C) or peak height (R = -0.21, p = 0.20, Fig. 3D) of the HRF. Further, there were no differences between platelet phenotype groups in AUC (F = 2.566, p = 0.09, Fig. 3F), TTP (F = 0.476, p = 0.63, Fig. 3G) or peak height (F = 2.364, p = 0.11, Fig. 3H).

The second principal component contributing to platelet reactivity (higher Sensitivity -> higher Capacity, PC2, 29.2% variance in platelet responses, Fig. 1C – 1E) was not associated with any measure of neurovascular function (FWHM, R = 0.26, p = 0.13; AUC, R = 0.12, p = 0.49; TTP, R = 0.29, p = 0.08; Peak Height, R = 0.12, p = 0.47).

Taken together, this establishes that higher platelet reactivity is associated with a shorter (HRF FWHM) and smaller (HRF AUC) cerebral blood flow response to activation of the visual cortex, with platelet reactivity multiparameter phenotyping revealing discernible impacts of platelet function on neurovascular function within healthy middle-age and older adults.

#### Evidence for mechanistic selectivity in the association between platelet reactivity and neurovascular function

Platelet regulation is complex and involves a balance between multiple activation and inhibition regulatory mechanisms, some of which overlap with signaling mechanisms governing neurovascular function. In particular, nitric oxide (NO) and prostanoids are key inhibitory regulators of platelet aggregation(*15*, *16*), and are simultaneously essential components of neurovascular function governing hemodynamic responses to neural activity(*14*). To determine whether specific aspects of platelet reactivity were associated with neurovascular function (HRF parameters), we analyzed individual platelet assay conditions with competing platelet activating and inhibiting manipulations. This provided evidence to suggest that there may be mechanistic selectivity that connects platelet reactivity and neurovascular function.

Specifically, higher platelet Sensitivity (EC50) and Capacity (Emax) were associated with shorter duration of the neurovascular response (HRF FWHM, Fig. 4A and 4B). This effect was observed most strongly when platelet aggregation was triggered by adenosine 5’-diphosphate (ADP) but was also present when platelet aggregation was triggered by Thrombin Receptor Activator Peptide 6 (TRAP-6). Platelet reactivity (Sensitivity or Capacity) was not associated with FWHM when platelet aggregation was triggered by Collagen-Related Peptide (CRP). This association between platelet reactivity and HRF FWHM was specific to the agonist used to trigger aggregation but was not affected by the presence or absence of competing platelet inhibition.

**Fig. 4.**
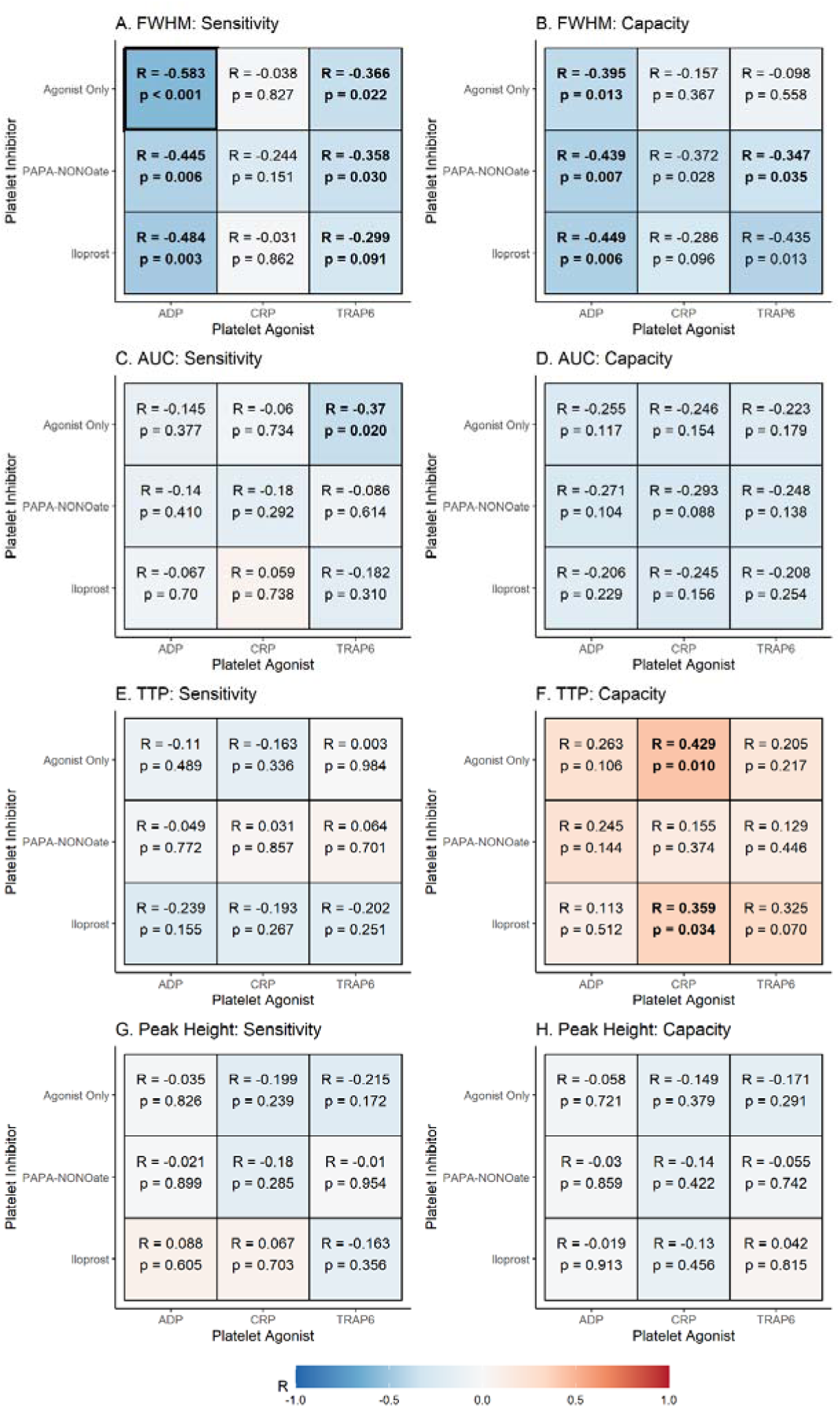
Evidence for mechanistic selectivity in the relationship between platelet reactivity and neurovascular function. (A) Relationships between platelet Sensitivity (EC50) and HRF full-width half-maximum (FWHM). (B) Relationships between platelet Capacity (Emax) and HRF FWHM. (C) Relationships between platelet Sensitivity and HRF area under the curve (AUC). (D) Relationships between platelet Capacity and HRF AUC. (E) Relationships between platelet Sensitivity and HRF time to peak (TTP). (F) Relationships between platelet Capacity and HRF TTP. (G) Relationships between platelet Sensitivity and HRF Peak height. (H) Relationships between platelet Capacity and HRF Peak height. All correlations are Spearman’s rho due to non-parametric data. Significant correlations (p < 0.05) are in bold text, while correlations surviving multiple comparison correction (False Discovery Rate; FDR) are displayed with bold borders. Scatter plots of each correlation is presented in Fig. S4.

Further, higher platelet Sensitivity (but not Capacity) was associated with an overall smaller neurovascular response (HRF AUC), but only when platelet aggregation was triggered by TRAP-6 (Fig. 4C). This association was specific to platelet reactivity to TRAP-6 without competing inhibition from iloprost (a synthetic prostanoid) or PAPA-NONOate (NO donor). Lastly, higher platelet Capacity (but not Sensitivity) was associated with a slower initiation of the neurovascular response (longer HRF TTP), but only when platelet aggregation was triggered by CRP (Fig. 4F). This association was specific to platelet responses to CRP, and persisted with the addition of competing inhibition by iloprost (a synthetic prostanoid). In contrast, the addition of PAPA-NONOate (NO donor) removed the association between platelet Capacity and HRF TTP.

No measure of platelet reactivity appeared to correlate with the maximum magnitude of the neurovascular response (HRF peak height, Fig. 4G and 4H).

Combined, the above findings provide evidence for mechanistic selectivity, such that specific platelet signaling mechanisms are associated with different physiological components of the hemodynamic response to neural activation. Though the functional implications of the observed effects are not fully understood, they are consistent with those observed in ageing(*19*), diabetes(*20*, *21*), and subjective cognitive decline(*22*), and are indicative of age-related attenuation in matching blood flow to energy demand in the active brain region.

#### The association between platelet reactivity and neurovascular function is not attributable to demographic differences

Partial correlations accounting for demographic variables of age, sex and BMI were conducted to determine whether the observed associations between platelet reactivity and neurovascular function simply reflected shared variance due to demographics (Fig. 5 and Table S1). Alternatively, additional relationships between platelet reactivity and neurovascular function could be obscured due to demographic effects.

**Fig. 5.**
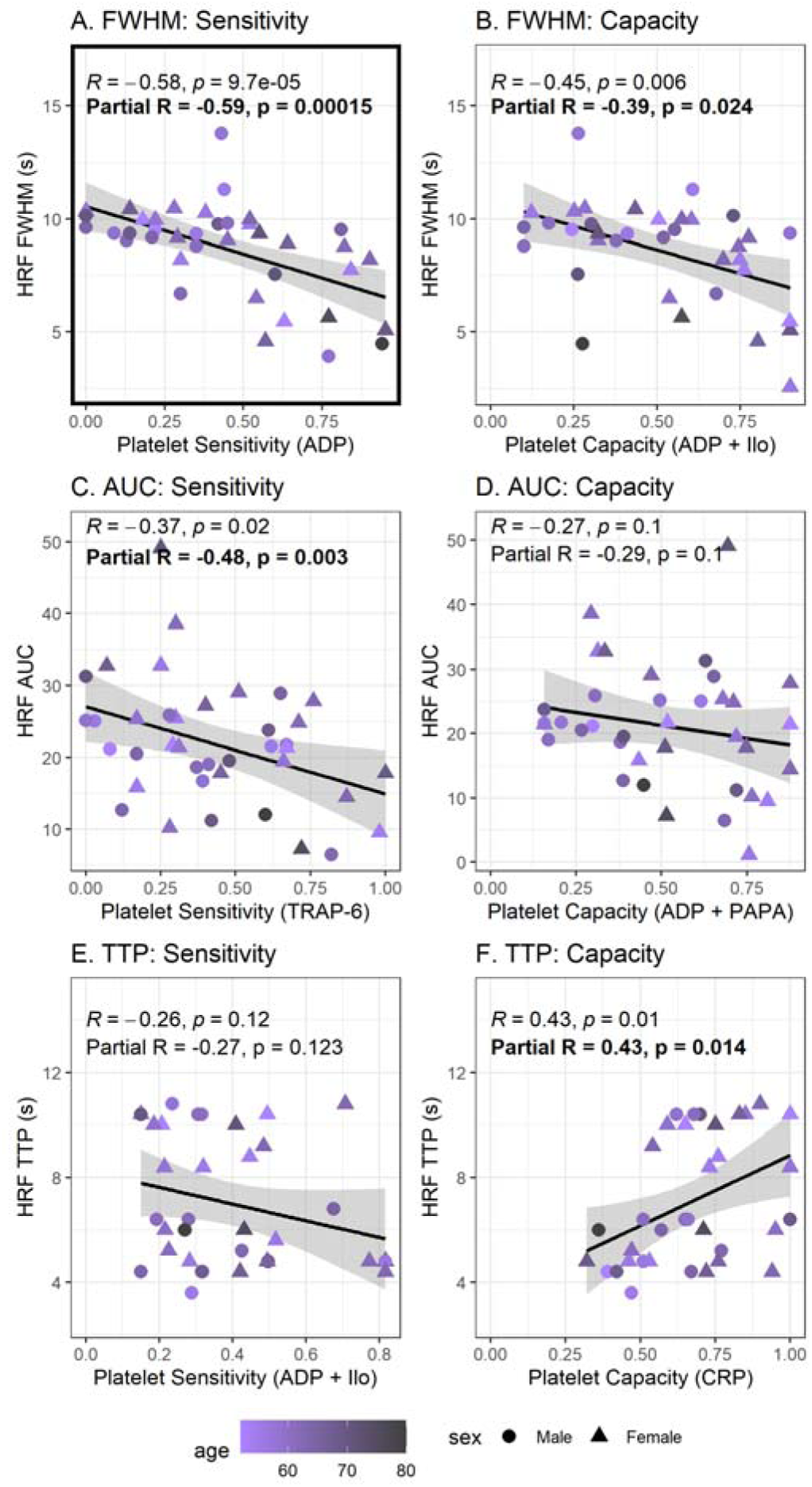
Relationships between platelet reactivity and neurovascular function are not attributable to demographic differences. The individual strongest correlation from each panel of Fig. 4 (A-F) is included here for illustrative purposes. Original correlations displayed in Fig. 4 were re-run accounting for demographic variables of age (color), sex (shape) and BMI (not shown on figure). Partial correlations between platelet reactivity and neurovascular function (hemodynamic response function (HRF) parameters) accounting for demographic variables (partial R) were not meaningfully different from the total correlations (R). (A) Relationship between platelet Sensitivity to Adenosine 5’-diphosphate (ADP; agonist only) and HRF full-width half-maximum (FWHM), (B) Relationship between platelet Capacity with ADP (+ iloprost) and HRF FWHM, (C) Relationship between platelet Sensitivity to thrombin receptor activator peptide 6 (TRAP-6) and HRF area under the curve (AUC), (D) Relationship between platelet Capacity with ADP (+ PAPA-NONOate) and HRF AUC, (E) Relationship between platelet Sensitivity to ADP (+ iloprost) and HRF time to peak (TTP), (F) Relationship between platelet Capacity with CRP (agonist only) and HRF TTP.

Our analyses suggested no direct influence of age or sex on neurovascular function (no significant partial correlations). Higher BMI was associated with smaller AUC and smaller peak height, suggesting high BMI is independently associated with smaller blood flow responses to neural activity.

Importantly, correlations between platelet reactivity and neurovascular function accounting for demographic variables were not meaningfully different from the total correlations (Fig. 5 and Table S1). This suggests that the association between platelet reactivity and neurovascular function does not simply reflect shared variance driven by demographic differences. In particular, the relationships between the multiparameter phenotyping-derived measure of latent platelet reactivity with shorter FWHM (total correlation R = -0.45, p = 0.006, Fig. 3A; partial correlation R = -0.46, p = 0.007) and smaller AUC (total correlation R = -0.41, p = 0.015, Fig. 3B; partial correlation R = -0.55, <0.001) were unchanged when accounting for age, BMI and sex. Platelet Sensitivity (ADP and TRAP-6) and Capacity (ADP) remained significantly correlated with HRF FWHM, platelet Sensitivity (TRAP-6) remained significantly correlated with HRF AUC, and platelet Capacity (CRP) remained significantly correlated with HRF TTP.

Two additional correlations became significant; (1) platelet Capacity with TRAP-6 + iloprost was significantly associated with longer HRF TTP after accounting for age, sex, and BMI (total correlation R = 0.33, p = 0.070; partial correlation R = 0.41, p = 0.026), (2) platelet Sensitivity to TRAP-6 was significantly associated with smaller HRF peak height after accounting for age, sex and BMI (total correlation R = -0.25, p = 0.134; partial correlation R = -0.37, p =0.027).

Therefore, although our data provides some indication that sex and BMI may affect neurovascular function as assessed by HRF parameter estimates, our observed associations between platelet reactivity and neurovascular function are not attributable to demographic factors.

#### Correction for multiple comparisons

The data included in this publication is part of a larger study which incorporates BOLD HRF parameters from other brain regions (outside V1; posterior cingulate cortex, precuneus, motor cortex) and cognitive tasks (episodic memory) obtained in the same scanning sessions. We chose to take the conservative approach to correct for multiple comparisons over all BOLD HRF parameters, platelet variables, and cognitive tasks included in the study (a total of 288 comparisons), including those not described in this publication.

Only one correlation remained after correction by false discovery rate (FDR) to an alpha of 0.05 (Fig. 4A, indicated by black box outline). Platelet Sensitivity to ADP (alone, without inhibitor) was negatively correlated with FWHM in V1 during visual stimulation (R = -0.58, FDR adjusted p = 0.028). This correlation also remained after FDR correction when accounting for demographic variables of age, sex, and BMI (R = -0.59, p = 0.0001, FDR adjusted p = 0.029).

Notably, ADP is the only agonist from the assay that not only triggers and potentiates aggregation but is also produced by platelets in vivo(*24*). This autocrine activation loop amplifies platelet aggregation and thrombus formation, underscoring the importance of platelet ADP responses in vascular physiology and pathology.

The correlation between platelet Sensitivity to ADP (alone; without inhibitor) and HRF FWHM was then interrogated to determine whether it was explained by systemic vascular function (combined peripheral and cerebral) or either peripheral vascular function or cerebrovascular function alone.

### Platelet reactivity directly affects neurovascular function

#### Systemic vascular function does not explain the association between platelet reactivity and neurovascular function

Given that dysregulation of the hemostatic system is a central aspect of cardiovascular disease pathophysiology(*8*), and concurrently, vascular dysregulation is a key contributor to dementia pathophysiology(*9*), we assessed whether the association between platelet reactivity and neurovascular function was explained through systemic vascular function. We used a serial mediation model to test whether the association between platelet reactivity (Sensitivity to ADP) and neurovascular function (HRF FWHM) was explained by a mechanistic pathway through peripheral (arm) and cerebral (brain) vascular function (schematic in Table 1). Peripheral vascular function was assessed by peripheral vascular reactivity to nitric oxide (NO), including change in perfusion in response to acetylcholine (ACh) which triggers NO release from the endothelium (endothelium-dependent), and in response to sodium nitroprusside (SNP) which is an NO donor that acts directly on the smooth muscle (endothelium-independent). Cerebrovascular function was assessed by cerebrovascular reactivity to carbon dioxide (CO_2_), including whole brain perfusion and arrival time under different CO_2_ conditions, and change in perfusion in response to hypercapnia (CVRhyper) and hypocapnia (CVRhypo) from the whole brain and specifically from the primary visual cortex (V1).

**Table 1.**
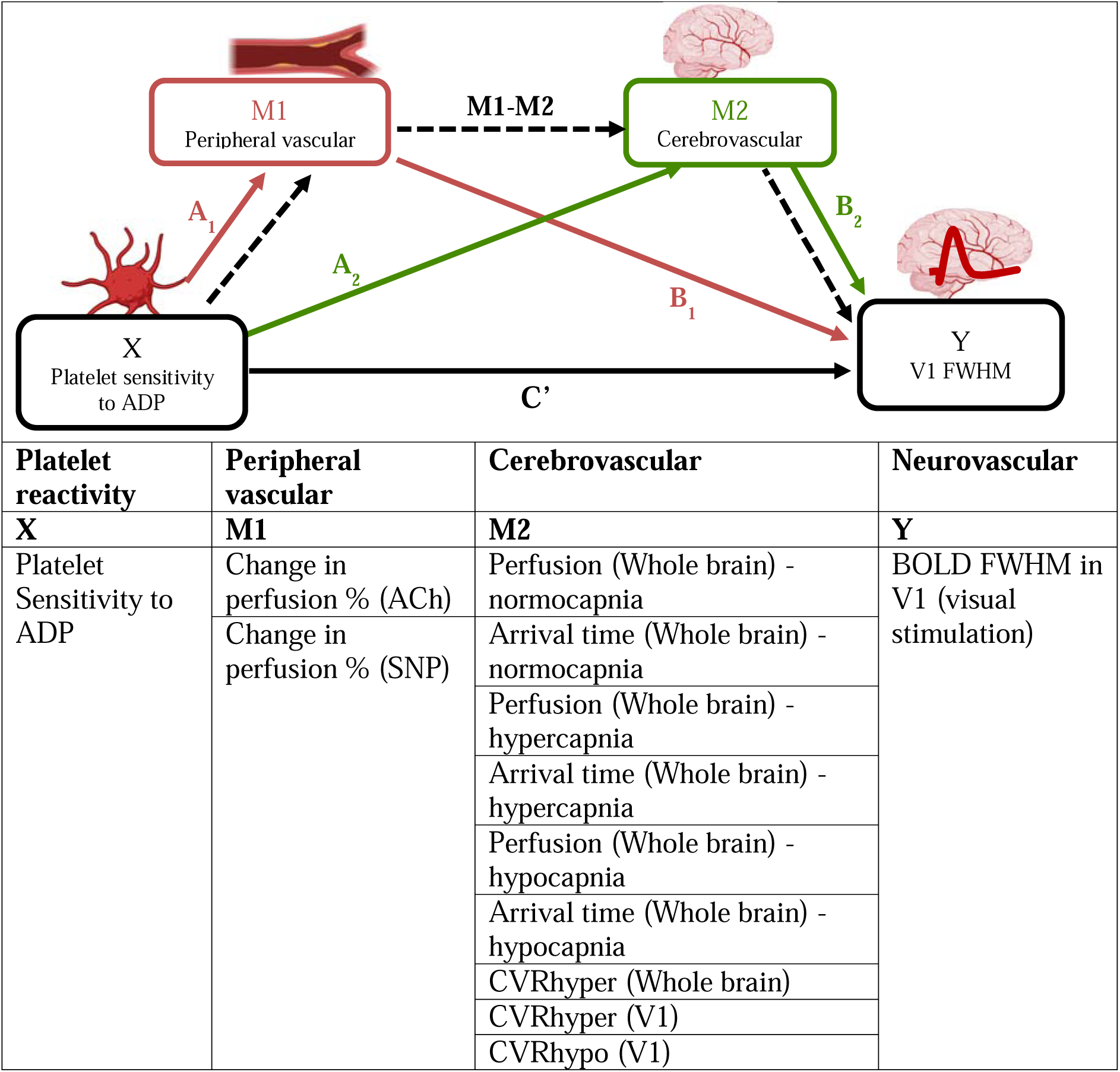
Schematic representation of serial mediation analyses including mediating variables. A total of 18 serial mediation models were tested to investigate whether the association surviving correction by false discovery rate (FDR) (between platelet Sensitivity to ADP and HRF FWHM) was explained by a serial pathway through peripheral vascular effects (M1) and cerebrovascular effects (M2). ACh, Acetylcholine; ADP, Adenosine 5’-diphosphate; BOLD, Blood Oxygen Level Dependent MRI signal; CVRhyper, Hypercapnic Cerebrovascular Reactivity; CVRhypo, Hypocapnic Cerebrovascular Reactivity; FWHM, Full-width Half Maximum; SNP, Sodium Nitroprusside (Nitric Oxide donor); V1, Primary Visual Cortex.

#### Systemic vascular function: Combined peripheral and cerebrovascular reactivity

The serial mediation model revealed that the association between platelet reactivity and neurovascular function was not explained by systemic vascular function determined by combined peripheral vascular reactivity and cerebrovascular reactivity. This suggests that platelets may directly interact with neurovascular function. No combination of peripheral and cerebral vascular function variables significantly explained the relationship between platelet Sensitivity to ADP and HRF FWHM (Table 2). Specifically, estimates of the indirect (mediating) effect ranged -0.001 to 0.019 (p values ranged 0.39 to 0.99) for all combinations of the vascular function variables.

**Table 2.**
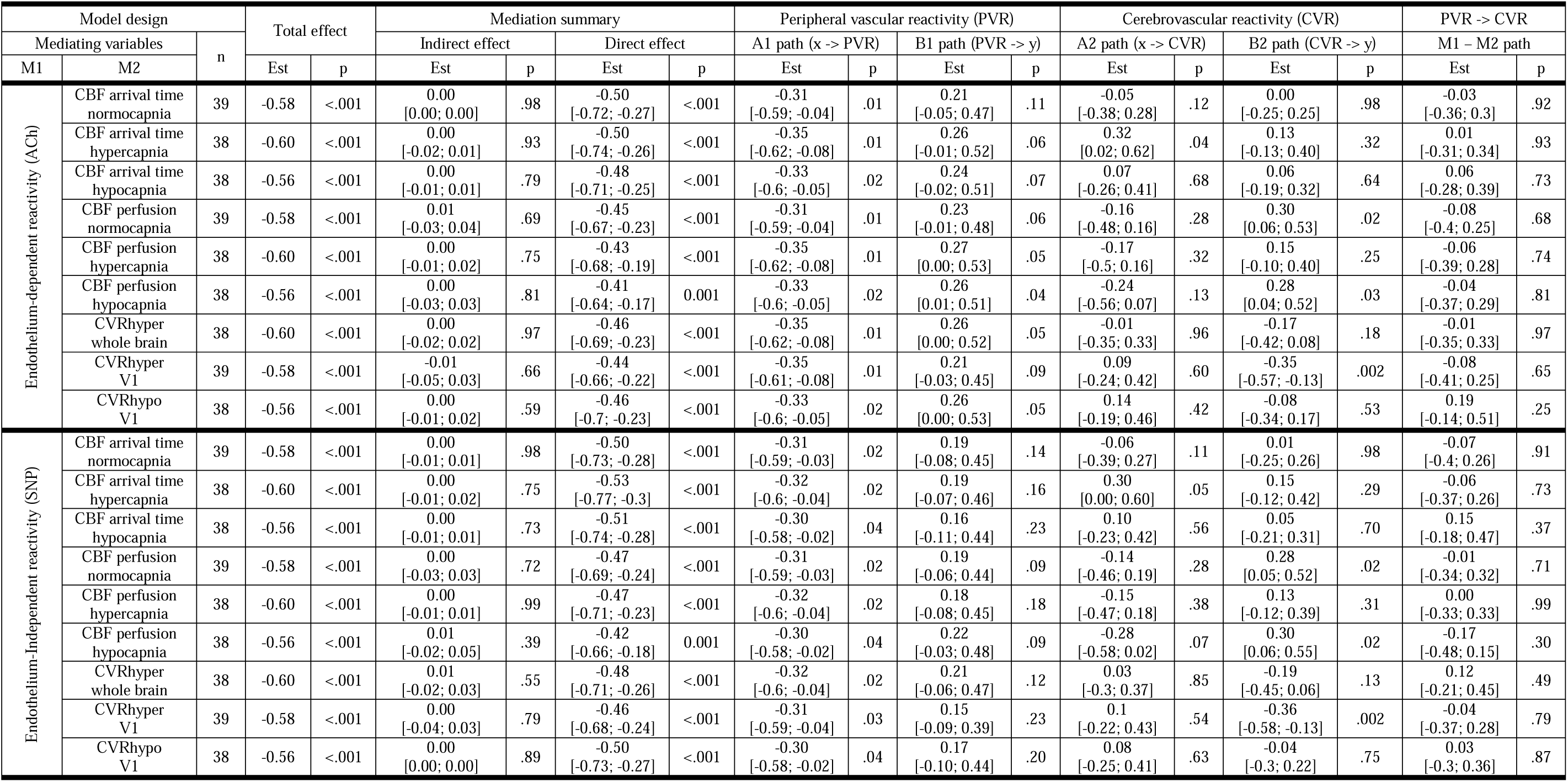
Systemic vascular function does not explain the relationship between platelets and neurovascular function: Serial mediation analyses. No combination of M1 and M2 variables significantly explained the relationship between platelet Sensitivity (x = Sensitivity to ADP) and the duration of the neurovascular response (y = HRF FWHM). ACh, Acetylcholine; CBF, Cerebral Blood Flow; CVRhyper, Hypercapnic Cerebrovascular Reactivity; CVRhypo, Hypocapnic Cerebrovascular Reactivity; Est, estimated effect; SNP, Sodium Nitroprusside (Nitric Oxide donor); V1, Primary Visual Corte

Of note, peripheral vascular reactivity was not associated with cerebrovascular reactivity, the estimate of the M1 -> M2 effect ranged -0.17 to 0.19 (p = 0.25 to 0.99).

#### Peripheral vascular reactivity to nitric oxide (NO)

The mediation analysis was also used to determine whether the association between platelet reactivity and neurovascular function (pathway C) was explained specifically by peripheral vascular reactivity to NO (excluding cerebrovascular reactivity). Platelet Sensitivity to ADP was associated with peripheral vascular reactivity to NO (pathway A1), but peripheral vascular reactivity was not associated with HRF FWHM (pathway B1), meaning peripheral vascular reactivity alone did not explain the association between platelet Sensitivity to ADP and HRF FWHM (Table 2).

Within the mediation model, the A1 and B1 pathways indicate whether the association between platelet Sensitivity to ADP and HRF FWHM was explained by peripheral vascular reactivity. Platelet Sensitivity to ADP had a moderate negative effect on both endothelium-dependent peripheral vascular reactivity (ACh) and endothelium-independent peripheral vascular reactivity (SNP) (A1 path estimates range -0.35 to -0.30; p = 0.01 to 0.04). This negative association likely reflects shared biochemical regulation of platelet inhibition and peripheral vascular tone regulation(*15*, *16*). However, peripheral vascular reactivity did not affect HRF FWHM in 17 out of the 18 mediation analyses (B1 path estimates range 0.15 to 0.27, p = 0.04 to 0.23). Combining these two paths (A1 -> B1) confirmed that the association between platelet Sensitivity to ADP and HRF FWHM was not explained by peripheral vascular reactivity (indirect effect estimates range -0.09 to -0.05, p = 0.12 to 0.30).

#### Cerebrovascular reactivity to carbon dioxide (CO_2_)

Finally, the mediation analysis was used to determine whether the association between platelet reactivity and neurovascular function (pathway C) was explained specifically by cerebrovascular reactivity to CO_2_ (excluding peripheral vascular reactivity). Although some measures of cerebrovascular function were associated with HRF FWHM (pathway B2), this was separate from the association with platelet Sensitivity to ADP (pathway C), meaning cerebrovascular function alone did not explain the association between platelet Sensitivity to ADP and visual FWHM (Table 2).

The A2 and B2 pathways of the mediation model indicate whether the association between platelet Sensitivity to ADP and HRF FWHM was explained by cerebrovascular function. Platelet Sensitivity to ADP had no effect on cerebrovascular function in 17 out of 18 mediation analyses (A2 path estimates ranged -0.35 to 0.30; p = 0.04 to 0.96). However, some cerebrovascular function variables were associated with HRF FWHM. CBF perfusion (normocapnia) was positively associated with HRF FWHM (B2 path estimate 0.28, p = 0.02), CBF perfusion (hypocapnia) was positively associated with HRF FWHM (B2 path ranged 0.28 to 0.30, p = 0.02) and finally, CVRhyper in the V1 was negatively associated with HRF FWHM (B2 path estimates -0.36 to 0.35, p = 0.002). Combining these two paths (A2 -> B2) confirmed that the association between platelet Sensitivity to ADP and HRF FWHM was not explained by cerebrovascular reactivity (indirect effect estimates ranged -0.09 to 0.04, p = 0.11 to 0.30).

## Discussion

This is the first study to establish a direct mechanistic link between platelet reactivity and neurovascular function in vivo in humans. Higher platelet reactivity was associated with a shorter duration and overall smaller hemodynamic response to activation of the primary visual cortex. This association captures a direct and specific link from platelet function to neurovascular coupling, one that bypasses systemic vascular function effects.

This study captured individual differences in neurovascular function in a sample of healthy middle-aged and older adults from the general population. Individuals with higher platelet reactivity exhibited attenuated (shorter, smaller, and delayed) hemodynamic responses to visual cortex activation, indicative of problems recruiting the vascular response to match the energy demand in active brain regions that may in-turn contribute to neurodegeneration. The hemodynamic response is pivotal for ensuring the delivery of energy substrates and removal of waste products from the brain(*25*, *26*). Failure to supply sufficient oxygenated blood can lead to hypoxic injury, potentially causing damage to neurons, impairing synaptic function, and ultimately resulting in brain atrophy(*27*, *28*). Chronic hypoxia and impaired clearance of metabolic waste products could induce neurotoxicity, triggering inflammation, oxidative stress, and causing damage to both neurons and glial cells(*14*). These neurotoxic processes significantly contribute to neurodegeneration(*29*). Chronically attenuated hemodynamic responses may initiate a cascade of detrimental events that contribute to the development and progression of dementia.

Though the functional implications of these novel findings are not yet fully understood, the observed effects in our study are consistent with previous literature demonstrating altered neurovascular coupling in clinical states. For example, subjective cognitive decline has been associated with shorter FWHM and longer TTP compared to age-matched controls(*22*). Older age has been associated with a longer TTP(*19*), while diabetes has also been associated with both a longer TTP and a smaller area under the curve (AUC)(*20*, *21*). These previous studies also reported smaller HRF peak heights compared to controls, but there were no associations between platelet reactivity and HRF peak height in the present study. We also found no effect of age on any HRF parameters, which contrasts previous work that specifically investigated age-effects in healthy adults(*19*). However, this discrepancy is reconciled by the fact that West and colleagues compared a group of younger adults (18 to 30 years) to a group of older adults (54 to 74 years), while the present study investigated a continuous effect of age within older adults only (50-80 years).

It is interesting that platelet reactivity to specific agonists was related to specific components of the hemodynamic response. Platelet aggregation in response to ADP was specifically related to narrower HRF FWHM, while platelet aggregation in response to thrombin receptor activator peptide 6 (TRAP-6) was specifically associated with smaller HRF AUC, and platelet aggregation in response to CRP was specifically related to slower TTP. ADP is released by activated platelets at the site of injury and potentiates platelet aggregation by binding to purinergic receptors on platelets, leading to a conformational change of the integrin αIIbβ3 on the platelet surface, allowing fibrinogen binding and platelet aggregation(*24*). This autocrine positive feedback loop amplifies platelet aggregation and thrombus formation. In contrast, TRAP-6 is a peptide that mimics the effects of thrombin and promotes platelet activation and aggregation via activation of protease-activated receptors on the platelet surface(*30*). CRP is a cross-linked peptide that mimics the action of collagen, which is exposed upon vascular injury. CRP (and collagen) initiates platelet adhesion and activation by binding to the glycoprotein VI (GPVI) receptor on platelets(*31*). These differing roles in the hemostasis cascade provide scope for future mechanistic studies to investigate, though it is worth noting that the only relationship remaining after correction for multiple comparisons was platelet sensitivity to ADP; the only agonist from the platelet assay that is released by platelets *in vivo*.

This study establishes a specific link between platelet reactivity and neurovascular function, independent of systemic vascular reactivity. The exact mechanisms are beyond the scope of this paper, but are hinted at by the multifaceted biological overlap between the two systems. For instance, serotonin release by activated platelets(*32*, *33*) may connect platelet activity to neurovascular function by influencing neurotransmission and vascular tone, acting on serotonin receptors present on neurons, glial cells, and vascular smooth muscle cells(*34*). Serotonin can cause vasoconstriction, potentially altering blood flow and neural activity(*35*). Over longer timescales, platelets may influence neurovascular function by affecting blood-brain barrier integrity and pericyte activity. Factors released by activated platelets, such as VEGF(*36*), TGF-β(*37*), and PDGF-BB(*38*), can increase BBB permeability, alter pericyte metabolism, and regulate vascular tone and immune responses.

Not only was the relationship between platelet and neurovascular function not explained by systemic vascular function, there was also no evidence for consistent ‘systemic vascular function’ (i.e. agreement between peripheral and cerebral vascular reactivity). This is in line with what others have observed(*39*), although the previous literature compared large blood vessels, whereas the present study specifically investigated microvascular perfusion in both the arm and the brain. This could be due to physiological differences between the peripheral vasculature and cerebrovasculature, or could simply be due to different vasoactive experimental manipulations (NO versus CO_2_).

Cerebrovascular function (assessed through cerebrovascular reactivity) and neurovascular function (assessed through HRF parameters) have shared physiological bases. However, to our knowledge, this is the first study to directly demonstrate a relationship between these measures in humans. HRF FWHM was positively associated with multiple (non-neuronal) cerebrovascular measures; cerebral blood flow (CBF) in normocapnia, CBF in hypocapnia, and cerebrovascular reactivity to hypercapnia (CVRhyper). It is particularly interesting that the duration of the hemodynamic response (HRF FWHM) was associated with the magnitude of the blood flow response to hypercapnia. While the initiation of the BOLD response is purported to relate to synaptically-driven feedforward signaling, the sustained BOLD response is purported to relate to feedback signaling driven more by CO_2_ and waste products(*40*). This finding therefore provides important empirical evidence to support this mechanistic picture, using integrated functional measures in humans.

This study is limited by its reliance on the BOLD response to infer neurovascular function. It is possible the reduced hemodynamic responses observed in this study may in fact reflect diminished underlying neural activity with intact neurovascular coupling since the BOLD signal is unable to differentiate between the neural and vascular components of neurovascular function. However, it is unclear through what mechanism higher platelet reactivity could affect neuronal activity isolated from neurovascular function. In contrast, as outlined above, there are plausible physiological mechanisms that could explain an association between platelet reactivity and neurovascular function. Another limitation of this study is the use of a cross-sectional design. A longitudinal design would have provided evidence for directionality, and insights on the functional implications of the observed relationships, allowing us to map the trajectory of potential neurodegeneration. Our findings indicate that such investigation would help not only further understand but also capitalize on the phenomenon observed herein. This paper is also limited by the focus on the visual cortex (V1) and did not include measures of cognitive function, or performance-based tasks, but the dataset produced by our broader project will allow us and others to address some of these questions.

In conclusion, we show that platelet reactivity was directly associated with neurovascular function in a healthy older sample. This novel direct association appears to bypass systemic vascular function, and to be mechanistically specific; distinct platelet signaling mechanisms were associated with different physiological components of the hemodynamic response to neural activity. Altered neurovascular function (hemodynamic responses to neural activation) marks the initial physiological stage in a cascade of events contributing to neurodegeneration. Therefore, understanding the precise mechanisms responsible for this novel finding may provide a substantial step forward in preventing this global health concern.

## Materials and Methods

### Study design

The study consisted of cross-sectional data collection that occurred over two study visits, separated by 7-31 days (mean (SD): 17.7 (8.9)). The *a priori* objectives of the study were to (i) determine whether platelet reactivity was associated with aspects of neurovascular function, and (ii) investigate whether specific aspects of platelet reactivity were associated with neurovascular function (mechanistic selectivity), and finally (iii) to test whether any association between platelet reactivity and neurovascular function could be explained by the state of the peripheral and cerebral vasculature. After initiation of the data analyses, we additionally sought to determine (iv) whether correlations between platelet reactivity and neurovascular function were directly attributable to platelet reactivity, or simply reflected shared variance driven by demographic differences.

Fifty-one healthy middle-aged and older adults (50-80 years; 28 females) were recruited into the study (mean (SD); age, 63.6 (7.7) years; BMI 25.5 (4.1) kg/m^2^). Eleven participants were excluded from the final analysis sample due to insufficient platelet or fMRI data for the primary analyses (final N = 40; 22 females; age, 61.9 (6.5) years; BMI 25.4 (4.1) kg/m^2^). Where other individual subjects were excluded from specific analyses, this is highlighted in the analysis description. Participants had no cardiovascular disease, diabetes, liver disease, hypertension, bleeding disorder, neuropsychiatric condition, current psychopathology, or sleep disturbance. Participants were not taking anticoagulant, antiplatelet, or psychoactive medications. Education level was 5 (2) across the sample and ranged from level 2 (e.g. GCSE grade A*-C) to level 8 (e.g. PhD).

### Ethical Approval

A favorable opinion for conduct was obtained from the University of Reading Research Ethics Committee (ref: UREC 21/11, accepted 14/06/2021) and the study was conducted following the standards of the World Medical Association Declaration of Helsinki 2013(*41*), with written informed consent obtained from all participants.

### Experimental procedures

For both visits, participants consumed a standardized meal the night before and fasted for 12 h before each study visit. Participants avoided strenuous exercise and alcohol and were advised to get a good night’s sleep the day before the visit. All study visits started at 08:00. Visit 1 included some procedures and measures not reported in this publication, but described in brief here for accuracy. Visit 1 consisted of a 29-minute encoding session for a subsequent memory task and a clinical assessment of memory function (not included in this publication), followed by a blood sample for platelet reactivity assaying, and MR Imaging of anatomical measures and functional MRI to measure task-evoked hemodynamic (BOLD) responses in the primary visual cortex (V1) during visual stimulation (and in multiple regions during an episodic memory task, not included in this publication). Visit 2 consisted of Laser Doppler Imaging (LDI) with iontophoresis of vasoactive agents to assess peripheral vascular reactivity followed by MR Imaging using arterial spin labelling (ASL) to measure CBF (reliability across visit 1 and visit 2: ICC = 0.76; R = 0.77; p < 0.001), hypercapnic CVR (5% CO_2_), and hypocapnic CVR (hyperventilation).

### Hemostatic function: Platelet reactivity

#### Sample collection

Blood samples were taken via venipuncture and collected into vacutainers containing 3.8% (w/v) sodium citrate. Platelet-rich plasma (PRP) was prepared by centrifuging whole blood at 100g for 20 mins and used within 30 mins.

#### Plate-based aggregometry (PBA) assay

Endpoint aggregation measurements were taken using a plate-based assay in PRP using 96-well half-area assay plates (Greiner) pre-prepared with freeze dried platelet agonists (collagen-related peptide (CRP), ADP and thrombin receptor activator peptide 6 (TRAP-6)) over a range of concentrations to cover the full concentration dose response for each (as described in Dunster et al(*23*)). PRP was first treated with vehicle, PAPA-NONOate (100µM), or iloprost (1.25nM) and incubated for 10 minutes at 37°C before the transfer of 40 µL to the assay plate. Plates were shaken at 1,200 rpm for 5 mins at 30°C using a plate shaker (Quantifoil Instruments) and absorption of 405 nm light measured using a FlexStation (Molecular Devices).

#### Assessment of platelet reactivity and multicomponent phenotyping

Data were analyzed using the in-house R package PPAnalysis to extract the metrics Sensitivity (EC50) and Capacity (Emax) and enable multiparameter phenotyping. A principal component analysis (PCA) was conducted on the vehicle data (platelet reactivity to activating agonists) using the prcomp function from the R package stats and clustered using an agglomerative hierarchical algorithm (as per Dunster et al(*23*)). This confirmed our previously published finding that platelet Sensitivity and platelet Capacity are distinct. This analysis provided individual assessment of latent platelet reactivity components for each participant, and phenotype stratification of platelet reactivity across the sample (Fig. 1). All platelet phenotypes were found to be stable assessed using the R package pvclust(*42*) with 1000 bootstrap iterations (Group 1, bp = 97%; Group 2, bp = 98%; Group 3, bp = 98%). Individual assay conditions (including competing platelet activating and inhibiting manipulations) were then interrogated to determine whether specific aspects of platelet reactivity were associated with neurovascular function.

### Neurovascular function

#### Visual Stimulation task

A black-and-white, radial checkerboard stimulus was presented to the participants to induce activation in the primary visual cortex (V1). The polarity of the checkerboard was reversed at a frequency of 8 Hz (every 125 ms), which has been shown to robustly activate V1 in older adults(*43*), and has been used in previous studies investigating neurovascular coupling in V1(*44*). The checkerboard stimuli were presented in 6.8 s blocks, with jittered inter-trial intervals (ITIs) between each period of visual stimulation which were optimized using optseq (Version 2.0, Center for Functional Neuroimaging Technologies, Massachusetts General Hospital, Harvard, US). During the ITIs, a blank grey screen with a white fixation cross in the center was displayed on the monitor (this was also presented for 2.72 s at the beginning of the task and 6.8 s at the end of the task). The total duration of the task was 310 s (5 min 10 s). The task was programmed using the psychtoolbox-3 extension of MATLAB(*45–47*) and was presented to the participants on a monitor with a resolution of 1024 × 768 and a refresh rate of 60 Hz.

#### MRI acquisition and analysis

Magnetic Resonance Imaging (MRI) data were collected on a SIEMENS Prisma-fit 3T scanner using a 32-channel head coil.

#### Anatomical reference: T1 MP-RAGE

High resolution T1-weighted images were acquired as an MP-RAGE sequence: time of echo (TE) = 2.29 ms, time of repetition (TR) = 2300 ms, time of inversion (TI) = 900 ms, Flip Angle (FA) = 8°, field of view (FOV) = 240 × 240 × 180 mm, voxel dimensions = 0.9 × 0.9 × 0.9 mm^3^, 192 slices (slice oversampling 16.7%), acquisition time = 5 min 21 s. T1 images were acquired for each scanning session and used for registration of the cerebral blood flow (CBF) and functional MRI (fMRI) scans to the MNI 152 2mm brain template using FMRIB’s Linear Image Registration Tool (FLIRT, FSL)(*48*).

#### Functional data: fMRI BOLD

To measure task-evoked activations, BOLD contrast functional images were acquired using an echo-planar (EPI) sequence (TR = 1360 ms, TE = 30 ms, FA = 90°, FOV = 192 × 192 × 100 mm^3^, voxel dimensions = 3 × 3 × 4 mm^3^, 25 slices). Echo-planar data quality were assessed for qualitative artefact (signal distortion e.g. ringing) and quantitative excessive motion (relative or absolute movement > 2mm) by agreement of two researchers assessing independently (GMKR and AS). The data were pre-processed in FSL including brain extraction (BET2), slice timing correction (interleaved), motion correction (MCFLIRT), spatial smoothing (FWHM = 5mm), high-pass filtering, B0 unwarping (fieldmap TR = 0.488 ms, TE1 =0.00519 ms, TE2 = 0.00765, FA = 60 °), and registration (FLIRT).

#### Estimation of hemodynamic response functions (HRFs)

We calculated individual hemodynamic response functions (HRFs) from pre-processed echo-planar data using a Finite Impulse Response (FIR) model in FSL with ten, 2-second impulses as regressors (onset from the start of each visual stimulation block, with a combined duration of 20 s). Individual statistical parametric maps for each impulse were then concatenated for each individual participant to create 4D niftis comprising of voxel-wise beta weights across the whole brain with each volume representing a 2-second impulse. A V1 region of interest (ROI) mask from the Harvard-Oxford subcortical atlas(*49*) (thresholded at 50% and binarized) was transformed into subject space for each participant, and used to extract the HRF time series for the V1 ROI of the visual stimulus regressor, creating a HRF that was specific to the participant, regressor, and ROI. HRF time series were then up-sampled to a frequency of 0.2 s and smoothed using a spline regression (R package = zoo(*50*)). Each individual HRF was qualitatively assessed by an experienced neuroscientist (GMKR) to determine whether it described physiologically-plausible task-evoked hemodynamics. HRF time series that were not physiologically plausible, or where the HRF could not be confidently identified, were excluded on the basis any extracted HRF parameters would reflect only noise (Fig. S2A). HRF parameters were calculated systematically but constrained by local minima corresponding to qualitative assessment of a single HRF impulse (Fig. S2B). This was necessary since the duration covered by HRF time series was set at 20 s across all cases, but clearly covered multiple HRF impulses in some cases. HRF parameters were calculated as described below (and shown in Fig 2D):

- Full-width half maximum (s) = width of the impulse at half of the peak height.
- Area under the curve = total quantity encompassed by the HRF impulse
- Time to peak (s) = time at which the peak height occurs.
- Peak height (%) = maximum BOLD signal value across the HRF impulse.

#### Experimental manipulation check: Visual stimulation evoked individualized hemodynamic response functions (HRFs)

An experimental manipulation check was conducted to ensure the visual stimulation task selectively activated the a priori ROI. Specifically, the visual stimulus regressor was associated with activations in V1. Group-level activation maps utilizing convolution with optimal basis functions (FMRIB’s Linear Optimal Basis Sets, FLOBS(*51*)) demonstrated clear activation localized to the visual cortex (Fig 2A), and recognizable HRFs were present in V1 during visual stimulation (Fig 2B). A large degree of individual variability existed regarding the shape of the HRF (Fig 2C), confirming the premise that task-evoked hemodynamics are not uniform, and may provide valuable information regarding individual neurovascular function. Individual parameters were then extracted from each HRF (Fig 2D).

Of note, BOLD HRF parameters were not correlated with each other. This suggests that at least in the context of short-duration stimulation (6.8 s) with a reversing visual checkerboard, the different HRF parameters capture different components of neurovascular function. Specifically, the BOLD FWHM did not correlate with TTP (R = 0.062; p = 0.7) or peak height (R = 0.17; p = 0.28), and the TTP did not correlate with peak height (R = 0.097; p = 0.53).

### Systemic vascular function

The maximum change in perfusion in response to a vasoactive stimulus is termed vascular reactivity, and is an important biomarker to assess vascular function. In this study vascular reactivity to vasoactive stimuli in the peripheral vasculature (arm) and cerebrovasculature (brain) was assessed to determine whether any associated between platelet reactivity and neurovascular function was explained by systemic vascular reactivity or cerebrovascular reactivity to vasoactive agents.

#### Peripheral vascular reactivity to nitric oxide (NO)

Peripheral vascular reactivity was assessed using Laser Doppler Imaging (LDI) with iontophoresis of vasoactive agents. The protocol is described in detail in Supplementary Methods. In brief, simultaneous delivery of acetylcholine (ACh, Sigma-Aldrich, Merck Ltd, Darmstadt, Germany) and sodium nitroprusside (SNP, Sigma-Aldrich, Merck Ltd, Darmstadt, Germany) was performed using an iontophoresis controller (MIC2, Moor Instruments, Devon, UK) to assess endothelium-dependent and endothelium-independent cutaneous perfusion, respectively. The results are presented as the percentage change in perfusion from the baseline to peak perfusion.

#### Cerebrovascular reactivity to carbon dioxide (CO_2_)

To measure cerebrovascular function, cerebral blood flow (CBF) measurements were acquired (i) at rest, (ii) in response to hypercapnia (5% CO_2_), and (iii) in response to hyperventilation-induced hypocapnia (30 breaths per minute(*52*)). Both hypercapnic CVR (CVRhyper) and hypocapnic CVR (CVRhypo) were calculated relative to individual change in end-tidal carbon dioxide (P_ET_CO_2_). Equations and detailed methods are provided in Supplementary methods. For all conditions, CBF was measured using arterial spin labelling (ASL). Images were acquired using the vessel-encoded PICORE Q2T pseudo-continuous ASL (pCASL) package. pCASL data are collected using an EPI readout (TR = 4000 ms, TE = 13.0 ms, TI = 1800 ms, FA = 90°, FOV = 220 × 220 × 118 mm^3^, voxel size = 3.4 × 3.4 × 4.5 mm^3^, 24 axial slices with 10% slice gap). Labelling of inflowing blood was achieved through a parallel slab applied below the acquisition slices (bolus duration = 700 ms, post-labelling delay (PLD) = 1800 ms post-labelling delay, 76 volumes, duration = 5 min 6 sec). Interleaved tag and control pairs were acquired with a tag RF flip angle (FA) of 20° and a duration of 600 µs, with 1000 µs separation. Perfusion and arrival time were quantified using the BASIL toolbox(*53*) within FSL (version 6.0.1).

### Statistical analysis

All statistical analyses were conducted in R (v 4.2.2). To determine whether platelet reactivity was associated with aspects of neurovascular function, Spearman’s rho correlations were conducted between principal component individual weightings and HRF parameters (full-width half maximum, FWHM (s); area under the curve, AUC; time to peak, TTP (s); peak height (%), peak). A 3 x 1 (phenotype group x HRF parameter) ANOVA was then conducted for each HRF parameter to compare the parameter between platelet phenotype groups. To investigate whether specific aspects of platelet reactivity (mechanistic selectivity) were associated with neurovascular function platelet reactivity, Spearman’s rho correlations were conducted between individual platelet assay conditions and HRF parameters. Correction for multiple comparisons by false discovery rate (FDR) was conducted across all correlations in the study, including those not reported in this article (FDR corrected alpha = 0.05). To determine whether correlations between platelet reactivity and neurovascular function simply reflected shared variance driven by demographic differences, partial correlations were run accounting for demographic variables of age, sex, and BMI. Finally, serial mediation models were conducted using the lavaan package(*54*) to test whether the association between platelet reactivity (Sensitivity to ADP) and neurovascular function (HRF FWHM) was explained by a mechanistic pathway through peripheral and cerebral vascular function. The total indirect effect (a1*d21*b2) was estimated through the model design, while the significance of individual mediator indirect effects was calculated using the distribution of the product of coefficients (Sobel Test).

## Acknowledgments

The authors would like to thank Shan Shen for her technical assistance.

## Author contributions

Conception and design of the work: GMKR, AC, JMG, JLD, JAL

Data Acquisition: GMKR, AS, BW, KMC, SR, EJ, EB

Data analysis: GMKR, JLD, AS, EJ

Writing – original draft: GMKR, JLD, JMG, AC

Writing – review & editing: GMKR, JLD, AS, BW, KMC, SR, EJ, EB, JAL, JMG, AC

## Data and materials availability

Pre-processed data is provided with this manuscript. Raw (skull-stripped) MRI data files are available on OpenNeuro (https://openneuro.org/).

## Supplementary Materials and Methods

### Peripheral vascular reactivity to Nitric Oxide (NO)

Both endothelium-dependent (acetylcholine, ACh) and endothelium-independent (sodium nitroprusside, SNP) peripheral microvascular reactivity were assessed using Laser Doppler Imaging (LDI, moorLDI2, Moor Instruments, Devon, UK) with iontophoresis. Simultaneous delivery of ACh (Sigma-Aldrich, Merck Ltd, Darmstadt, Germany) and SNP (Sigma-Aldrich, Merck Ltd, Darmstadt, Germany) was performed using an iontophoresis controller (MIC2, Moor Instruments, Devon, UK) to assess endothelium-dependent and endothelium-independent cutaneous perfusion, respectively. Perfusion changes in response to the delivery of both vasoactive drugs were assessed on the participant’s volar aspect of the right forearm. Following two basal measurements of skin perfusion, an incremental constant current was delivered using the LDI software. Current delivery was progressively increased in 5 mA steps (5, 10, 15, and 20 mA) to yield a total charge of 8 millicoulombs within the first 12 minutes, with 5 additional scans at the end delivering no current (0 µA) and therefore without the delivery of the vasoactive agents. A total of 21 scans were performed. ACh and SNP were diluted to 1% solutions with 0.5% saline and delivered simultaneously into the skin *via* anode (ACh) and cathode (SNP) internal electrode Perspex chambers (IZl22mm) (ION 6, Moor Instruments, Devon, UK). The scans were performed simultaneously with the iontophoresis protocol. The scan protocol lasted 21 min and all scans were performed with ambient lighting restricted.

Measurements of perfusion were conducted offline using the moorLDI Review V6.1 software. Perfusion values were quantified for ACh and SNP calculating the median for each region of interest. Results are presented as the percentage change in perfusion from the baseline scan collected immediately before the drug delivery, and was calculated as follows:

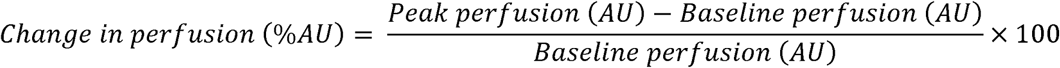

### Cerebrovascular Reactivity to carbon dioxide (CO2)

#### Hypercapnic cerebrovascular reactivity (CVRhyper)

Hypercapnic CVR (CVRhyper) was calculated relative to individual change in end-tidal carbon dioxide (P_ET_CO_2_), using the following equations:

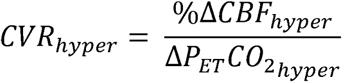

Where

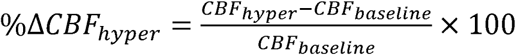

And

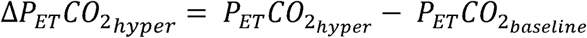

#### Hypocapnic cerebrovascular reactivity (CVRhypo)

To assess the decrease in vasodilation in response to a vasoconstricting stimulus, participants hyperventilated to induce hypocapnia for 5 min while a 5 min pcASL scan was acquired using the same parameters as for the baseline rCBF measurement detailed above. Hyperventilation of 30 breaths per minute was achieved through participants breathing in time to a visual cue. Based on previous literature this degree of hyperventilation should induce ∼20 mmHg decrease in end-tidal CO_2_. Hypocapnic CVR (CVR_hypo_) was calculated using the following equations:

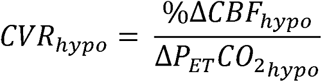

Where

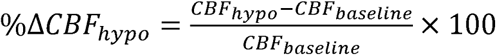

And

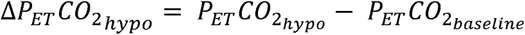

**Fig. S1.**
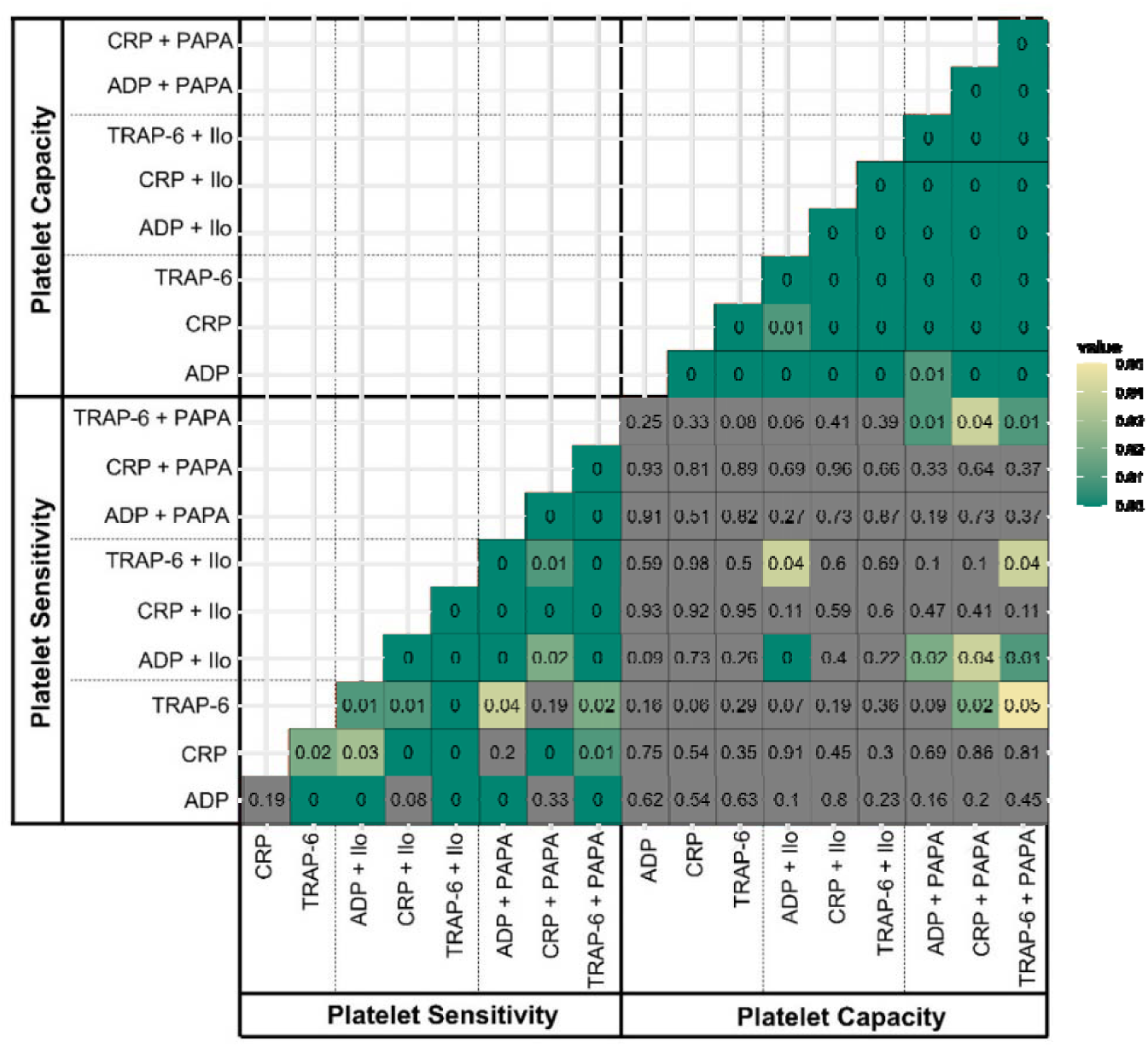
Matrix of p values within platelet reactivity metrics. Corresponding to Spearman’s rho values presented in Figure 1B of main text.

**Fig. S2.**
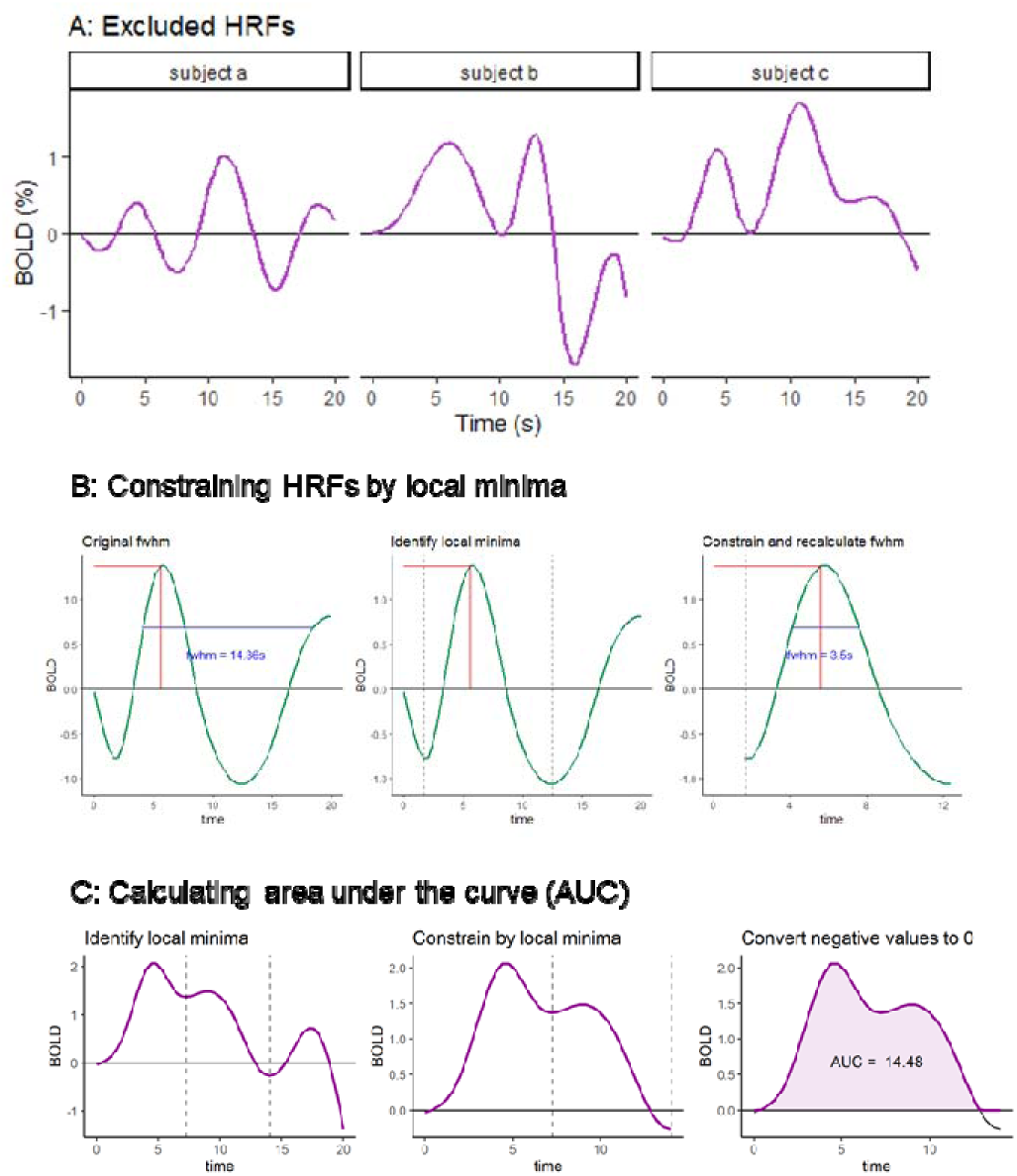
Schematic representation of qualitative assessment procedure for hemodynamic response functions (HRFs). (A) HRF time series that were not physiologically plausible, or where the HRF could not be confidently identified, were excluded on the basis any extracted HRF parameters would reflect only noise. (B) HRF parameters were calculated systematically but constrained by local minima corresponding to qualitative assessment of a single HRF impulse. This was necessary since the duration covered by HRF time series was set at 20s across all cases, but clearly covered multiple HRF impulses in some cases.

**Fig. S3.**
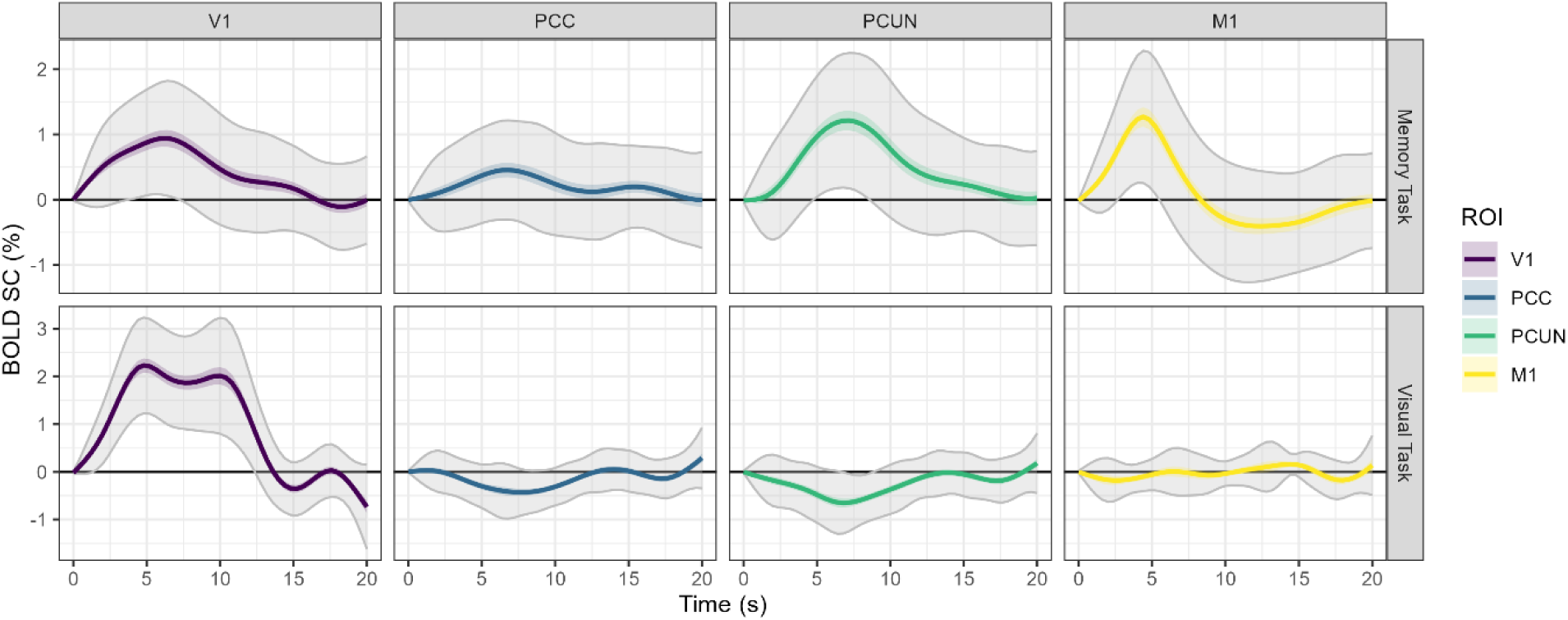
Group-averaged task-evoked hemodynamic response functions (HRFs) from multiple tasks and brain regions. Solid line values are the group mean, colored ribbons are ± standard error, and grey ribbons are ± standard deviation. Data from all participants is included in these figures, including hemodynamic traces from individuals with non-physiologically-plausible HRFs which were excluded from subsequent HRF parameter analyses. Group-level HRFs were present only in *a priori* task-relevant regions of interest (ROIs). M1, Motor Cortex (left, digit 1); PCC, Posterior Cingulate Cortex; PCUN, Precuneus; V1, Primary Visual Cortex.

**Fig. S4.**
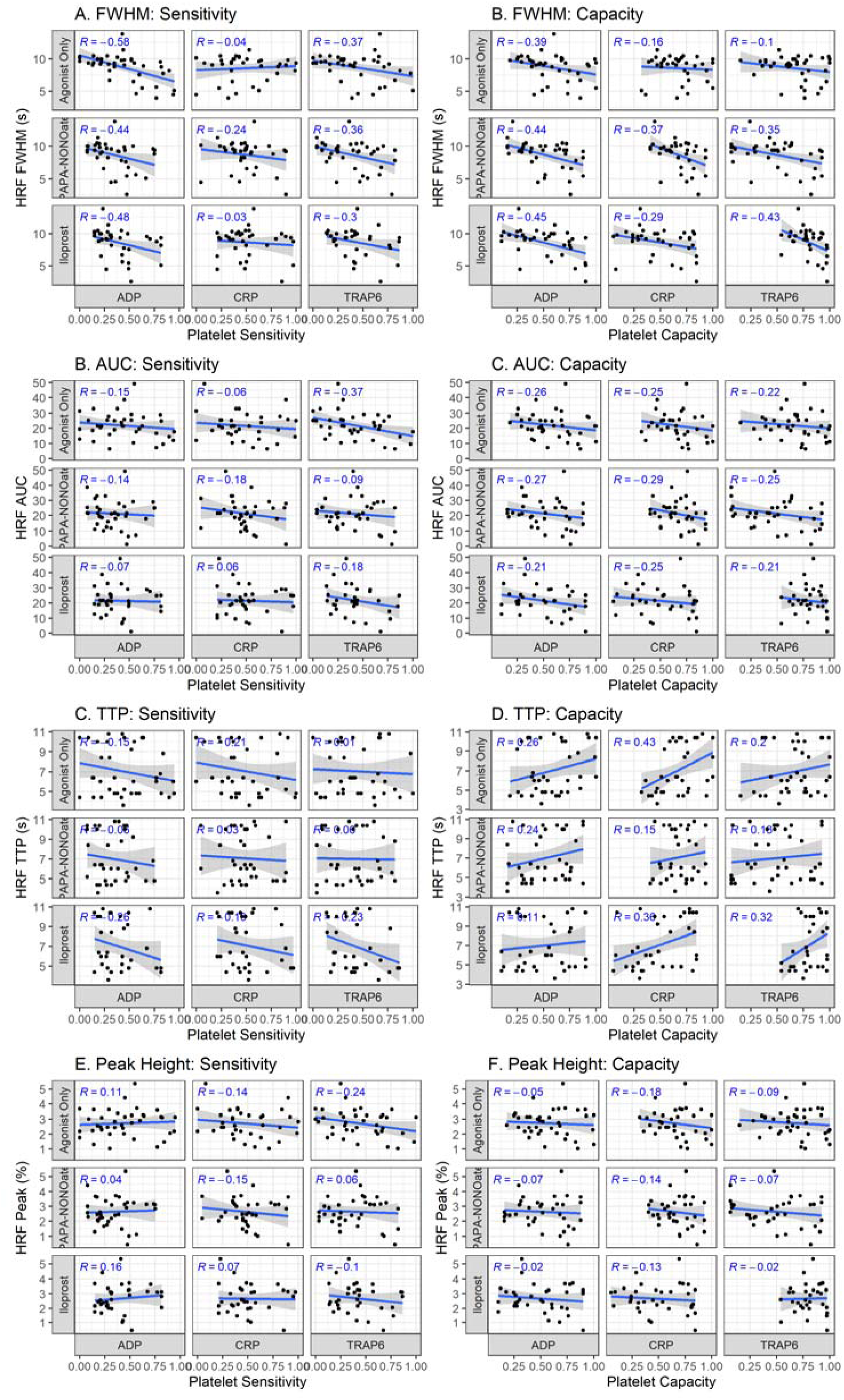
Scatter plots for Figure 4. All correlations presented as heatmaps in Fig 4 in the main text are presented as full scatter plots displaying the individual data points here.

## Table S1. (separate file)

Full results of partial correlation analyses. Corresponding to results section ‘The association between platelet reactivity and neurovascular function is not attributable to demographic differences’ and Figure 5. Correlations between platelet reactivity and neurovascular function accounting for demographic variables were not meaningfully different from the total correlations. Provided as a separate file in .xlsx format as it is a large table that extends beyond the page dimensions.

## Data S1. (separate file)

Preprocessed data for all variables included in statistical analyses.

